# The BAF chromatin complex component SMARCC1 does not mediate GLI transcriptional repression of Hedgehog target genes in limb buds

**DOI:** 10.1101/2023.02.03.527038

**Authors:** Janani Ramachandran, Wanlu Chen, Rachel K. Lex, Kathryn E. Windsor, Hyunji Lee, Tingchang Wang, Weiqiang Zhou, Hongkai Ji, Steven A. Vokes

## Abstract

Transcriptional responses to the Hedgehog (HH) signaling pathway are primarily modulated by GLI repression in the mouse limb. Previous studies suggested a role for the BAF chromatin remodeling complex in mediating GLI repression. Consistent with this possibility, the core BAF complex protein SMARCC1 is present at most active limb enhancers including the majority of GLI enhancers. However, in contrast to GLI repression which reduces chromatin accessibility, SMARCC1 maintains chromatin accessibility at most enhancers, including those bound by GLI. Moreover, SMARCC1 binding at GLI-regulated enhancers occurs independently of GLI3. Consistent with previous studies, some individual GLI target genes are mis-regulated in *Smarcc1* conditional knockouts, though most GLI target genes are unaffected. Moreover, SMARCC1 is not necessary for mediating constitutive GLI repression in HH mutant limb buds. We conclude that SMARCC1 does not mediate GLI3 repression, which we propose utilizes alternative chromatin remodeling complexes.

## INTRODUCTION

The Hedgehog (HH) pathway is a multifaceted regulator of embryonic development with distinct roles in regulating the formation of most organs. Diverse cell types respond to HH signaling through cell-specific transcriptional responses mediated by GLI transcription factors. GLI proteins are bifunctional, acting as context-dependent transcriptional activators (when HH signaling is active) or transcriptional repressors (in the absence of HH). GLI repressors are particularly important in the limb bud where most HH target genes are activated solely by a loss of GLI3, the primary GLI repressor (GLI de-repression) (Lewandowski et al., 2015; Litingtung et al., 2002; teWelscher et al., 2002). Constitutive GLI repression causes reductions in H3K27 acetylation (H3K27ac), a marker associated with active enhancers, as well as reductions in ATAC-seq accessibility (Creyghton et al., 2010; Heintzman et al., 2009; Lex et al., 2020; Rada-Iglesias et al., 2011). This suggests that GLI represses transcription by inactivating enhancers through interactions with an HDAC-containing repression complex. Additional unknown factors likely contribute to regulating chromatin accessibility as well as enhancer specificity (Lex et al., 2020). Several co-factors or complexes have been proposed to regulate GLI transcription based on physical and genetic association (Chen et al., 2004; Dai et al., 2002; Huang et al., 2016; Zhang et al., 2013, 2013). However, it has been challenging to specifically link them to GLI repression through standard genetic analysis since most co-factors are inherently pleiotropic, affecting the transcriptional regulation of multiple signaling pathways.

The SWI/SNF BAF complex has emerged as a top candidate for mediating both GLI activation and repression. This is based on evidence identifying physical interactions between multiple GLI proteins and several BAF complex members, mutations in various BAF components that alter expression of HH target genes, and that several BAF components bind to the promoters of HH target genes *Gli1* and *Ptch1* (Jagani et al., 2010; Jeon and Seong, 2016; Shi et al., 2016, 2014; Xiong et al., 2013; Zhan et al., 2011). BAF complexes are large, variable complexes that contain either SMARCA2 or SMARCA4, proteins with ATPase domains that mediate changes in nucleosomal density at enhancers or promoters. BAF complexes also include several core components, including SMARCC1 which promotes ATPase activity as well as the stability of BAF proteins (Clapier et al., 2017; Jagani et al., 2010; Sohn et al., 2007; Sokpor et al., 2017).

Since BAF complexes broadly maintain enhancer accessibility and H3K27ac enrichment across many different pathways, it is unclear how they might mechanistically mediate GLI repression (Alver et al., 2017; Hendy et al., 2022; Wang et al., 2017). As GLI transcriptional repression promotes reduced enhancer accessibility and H3K27ac depletion, it would seem more likely that these changes would be facilitated by loss of BAF complex activity rather than its presence. Additionally, the relatively global requirements for BAF complexes at many developmental enhancers indicate that they have upstream roles in regulating enhancer accessibility that might be distinct from GLI3 repression. Nonetheless, BAF complexes bind to GLI3 repressor isoforms and a loss of BAF activity results in the expansion of some GLI target genes (Jagani et al., 2010; Jeon and Seong, 2016; Zhan et al., 2011). Moreover, it has been reported that the BAF component SMARCA4 mediates both GLI activation and repression (Zhan et al., 2011). These studies, which examined a few signature target genes of the HH pathway are consistent with the notion that BAF complex could mediate GLI repression. In keeping with this possibility, BAF complexes can repress transcription in certain cases where they reposition a repressive nucleosome (Rafati et al., 2011). Additional scenarios in which GLI3 repressor complexes bind to and attenuate BAF complex activity are also possible.

To clarify the role of BAF complexes in HH gene regulation, we tested the hypothesis that SMARCC1 mediates GLI repression in limb buds. Consistent with previous reports, *Smarcc1* conditional mutants have reduced BAF complex stability (Choi et al., 2012; Sohn et al., 2007). Thousands of enhancers, including most GLI enhancers that are bound by SMARCC1 have markedly reduced chromatin accessibility in the absence of SMARCC1, suggesting that SMARCC1 is likely involved in enhancer activation rather than GLI-mediated repression, which is associated with reduced chromatin accessibility. We also report that GLI3 does not recruit or maintain SMARCC1 at most GLI enhancers. While some HH target genes are mis-regulated in SMARCC1 limb buds, their expression changes are generally not concordant with GLI repression. Finally, conditional deletion of *Smarcc1* is not sufficient to rescue the phenotypes caused by constitutive GLI repression in *Smarcc1*;*Shh^-/-^* double mutant limb buds. We conclude that the BAF component SMARCC1 does not mediate GLI repression in limb buds.

## RESULTS AND DISCUSSION

### SMARCC1 binds to most active limb enhancers and the majority of GLI-binding regions

To determine if BAF complexes are present at GLI enhancers in the mouse limb bud, we identified the genomic binding regions for SMARCC1, a core component of BAF complexes that serves a scaffolding role and protects other members of the complex from proteolytic degradation (Chen and Archer, 2005; Sohn et al., 2007). We dissected limb buds into anterior and posterior halves and identified a combined total of 48,268 SMARCC1-bound regions at E11.5 using CUT&RUN (Figure 1A). The majority of SMARCC1-bound regions (22,344) were common to both the anterior and posterior limb buds, and most (∼69%) were enriched for the active enhancer mark H3K27ac consistent with prior reports of BAF complex recruitment to H3K27ac-enriched active enhancers (Figure 1A-B; Source data 1) (Creyghton et al., 2010; Heintzman et al., 2009; Malkmus et al., 2021; Rada-Iglesias et al., 2011). To understand the extent to which SMARCC1 broadly regulates enhancers, we assessed SMARCC1 binding at H3K27ac ChIP-seq peaks as well as validated enhancers from the VISTA Enhancer Browser (Visel et al., 2007). Most SMARCC1-bound regions (79% of anterior and 71% of posterior regions) were enriched for H3K27ac (Supp. Figure 1A). Similarly, half of all H3K27ac-enriched regions (52%), and most (87%) of the annotated VISTA limb enhancers were bound by SMARCC1 (Figure 1B, Supp. Figure 1B-D), (Alver et al., 2017; Park et al., 2021). In addition to regions that were bound by SMARCC1 in both anterior and posterior limb bud halves, there were a smaller portion of anterior-specific (5,738) and posterior-specific (20,186) regions (Figure 1A). These included VISTA limb enhancers with enhancer activity in the same anterior/posterior region, suggesting that they correspond to spatially restricted limb enhancers (Figure 1C, Supp Figure 1B-D). These findings indicate that SMARCC1 is bound to many active enhancers.

**Figure 1.**
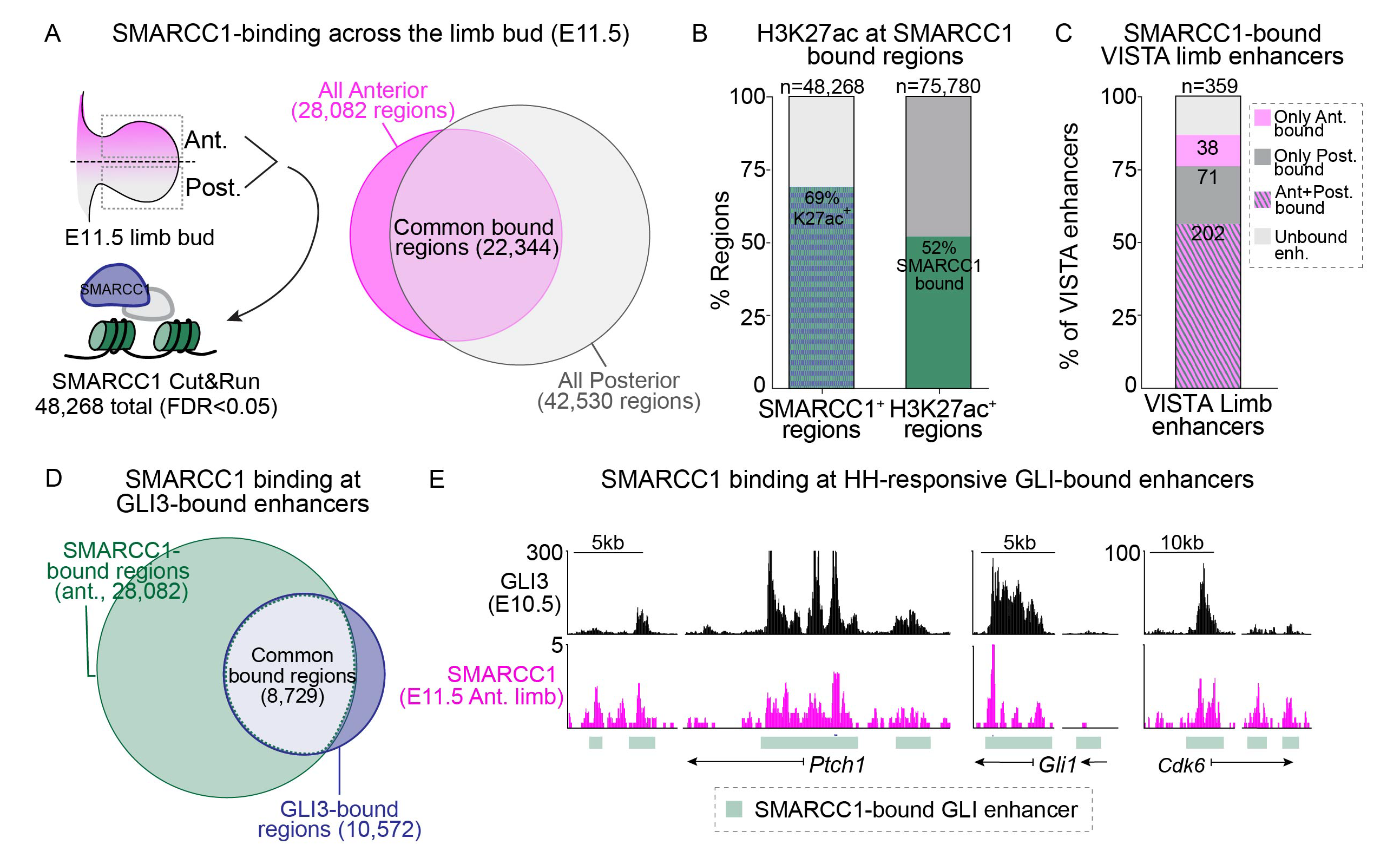
SMARCC1 binds to active limb enhancers and GLI binding regions. A. SMARCC1 binding regions in anterior and posterior halves of E11.5 wild-type forelimbs identified by CUT&RUN (n=3). B. SMARCC1 and previously identified H3K27ac binding in E11.5 forelimbs are co-enriched. C. Most VISTA limb enhancers are bound by SMARCC1. D. Intersection of all anterior SMARCC1-binding regions with previously identified E10.5 GLI3 binding regions. E. GLI3 anterior SMARCC1 binding at select GLI target genes.

To determine if BAF complexes are associated with GLI repressor-bound regions, we intersected SMARCC1 binding regions in anterior halves (where there are high levels of GLI repression) with a set of previously identified regions that are primarily or exclusively bound by GLI3-repressor (Lex et al., 2022). Most GLI3 binding regions (∼80%) were also bound by SMARCC1 in the anterior limb (Figure 1D-E), including the subset of HH responsive GLI binding regions (hereafter referred to as ‘GLI-enhancers’) that are enriched around HH target genes (Supp. Figure 1E) (Lex et al., 2020). We conclude that SMARCC1 binds to a large number of active enhancers in the limb, including regions bound by GLI3. We next sought to determine if SMARCC1 is required for regulating chromatin accessibility in response to GLI repression.

### SMARCC1 maintains chromatin accessibility throughout the genome including GLI enhancers

Since GLI repression is associated with a reduction in chromatin accessibility (Lex et al., 2022, 2020), we sought to determine if BAF complexes mediate this aspect of GLI repression by conditionally deleting *Smarcc1* in limb buds with *PrxCre* (here after referred to as ‘SMARCC1 cKOs’). Confirming previous findings, SMARCA4levels were reduced along with substantial reductions in most additional BAF complex members tested with the exception of SMARCC2 (Tuoc et al., 2013) (Supp. Figure 2A) (Jeon and Seong, 2016). Although SMARCC1 levels were also substantially reduced, they were not eliminated, suggesting the likelihood of some retained BAF complex activity. We then identified ATAC-accessible regions in the anterior halves of limb buds from E11.5 control and SMARCC1 cKOs (Figure 2A). More than 10% of all ATAC-accessible regions (10,906/100,188) had significantly reduced accessibility in SMARCC1 cKO limb buds (FDR<0.05), 58% (6,279/10,906) of which were enriched for H3K27ac (Figure 2B-D, E, Source data 3). Approximately half of GLI enhancers (154/309) had significantly reduced ATAC-accessibility in the absence of *Smarcc1*. Although not statistically significant, most of the additional GLI enhancers appeared less accessible in SMARCC1 cKOs. In total, 303/309 regions appear less accessible while 6 regions had a non-significant increase in accessibility (Figure 2B-D, E, Source data 3). Contrary to our initial predictions, only 62 out of all 10,906 significantly ATAC-changed regions and none of the 154 significantly ATAC-changed GLI-enhancer regions (Source data 3), gained accessibility in SMARCC1 cKOs, suggesting that rather than mediating GLI3-regulated chromatin compaction as a co-repressor, SMARCC1 has a major genome-wide role in maintaining enhancer accessibility.

**Figure 2.**
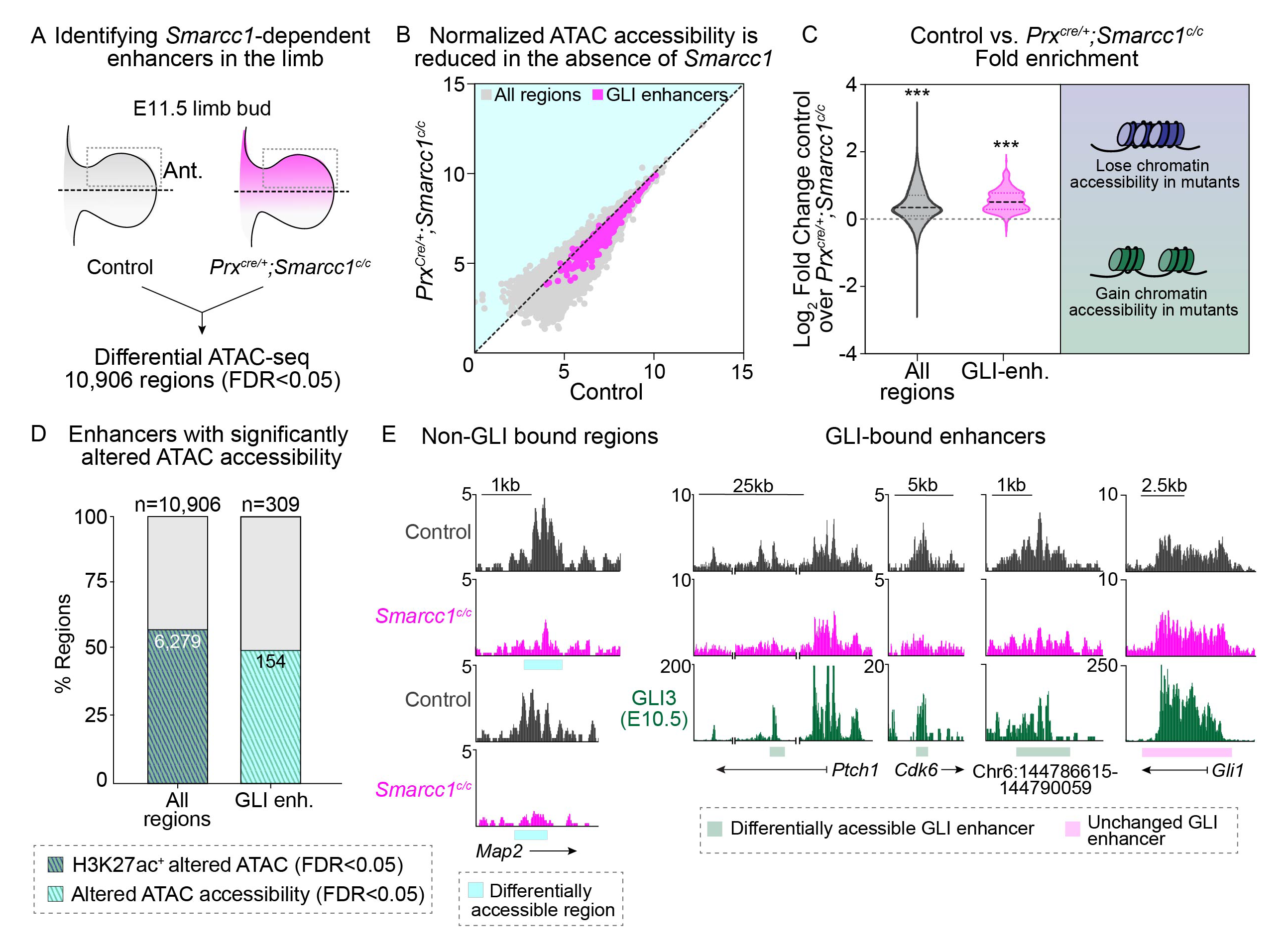
SMARCC1 has GLI-independent roles in regulating chromatin accessibility. A, B. Differential ATAC-seq on E11.5 control (*PrxCre^-^;Smarcc1^c/c^*; n=2) and *PrxCre^+^;Smarcc1^c/c^* (n=3) anterior forelimbs. Normalized enrichment plot indicates ATAC enrichment for all regions (grey; n=100,188) and all GLI enhancers (magenta; n=309). C. ATAC fold enrichment in control vs. *PrxCre^+^;Smarcc1^c/c^* samples across all ATAC accessible regions (grey) and GLI enhancers (magenta) showing significantly decreased accessibility in mutants (Wilcoxon signed rank test; ***p<2.2e-16). D. Approximately half of GLI enhancers have altered ATAC accessibility in *PrxCre^+^;Smarcc1^c/c^* forelimbs. E. Tracks showing GLI3 binding (green), and ATAC accessibility in control and mutant samples.

### GLI3 and SMARCC1 are independently recruited to nascent and active GLI enhancers

We asked if GLI3 might recruit SMARCC1 to nascent GLI enhancers by examining SMARCC1- binding in early limb buds prior to the apparent onset of GLI repression (Lex et al., 2022). At this stage, 60% (128/216; Source data 1) of the GLI enhancers that were bound by GLI3 in early limb buds were not bound by SMARCC1. Conversely, most GLI enhancers bound by SMARCC1 at E9.25 also had GLI3 enrichment at this stage (88/104), were enriched for H3K27ac (70/104), and positioned intragenically or proximal to gene promoters (82/104) (Figure 3A). These included *Gli1,* a HH-target gene that is primarily regulated by a promoter-proximal enhancer, and *Cdk6* (Figure 3 A-C) (Lopez-Rios et al., 2012; Vokes et al., 2008). This indicates that GLI3 binds most nascent GLI enhancers without SMARCC1, which becomes associated with most GLI enhancers only at later stages (Fig. 1D-E; Supp. Figure 1D). This is in contrast to previous reports that have shown BAF recruitment to poised early enhancers in human embryonic stem cells (Rada-Iglesias et al 2011) and suggests tissue-specific recruitment of BAF complexes to early limb bud enhancers. We conclude that GLI3 alone is not sufficient to recruit SMARCC1 to nascent limb enhancers.

**Figure 3.**
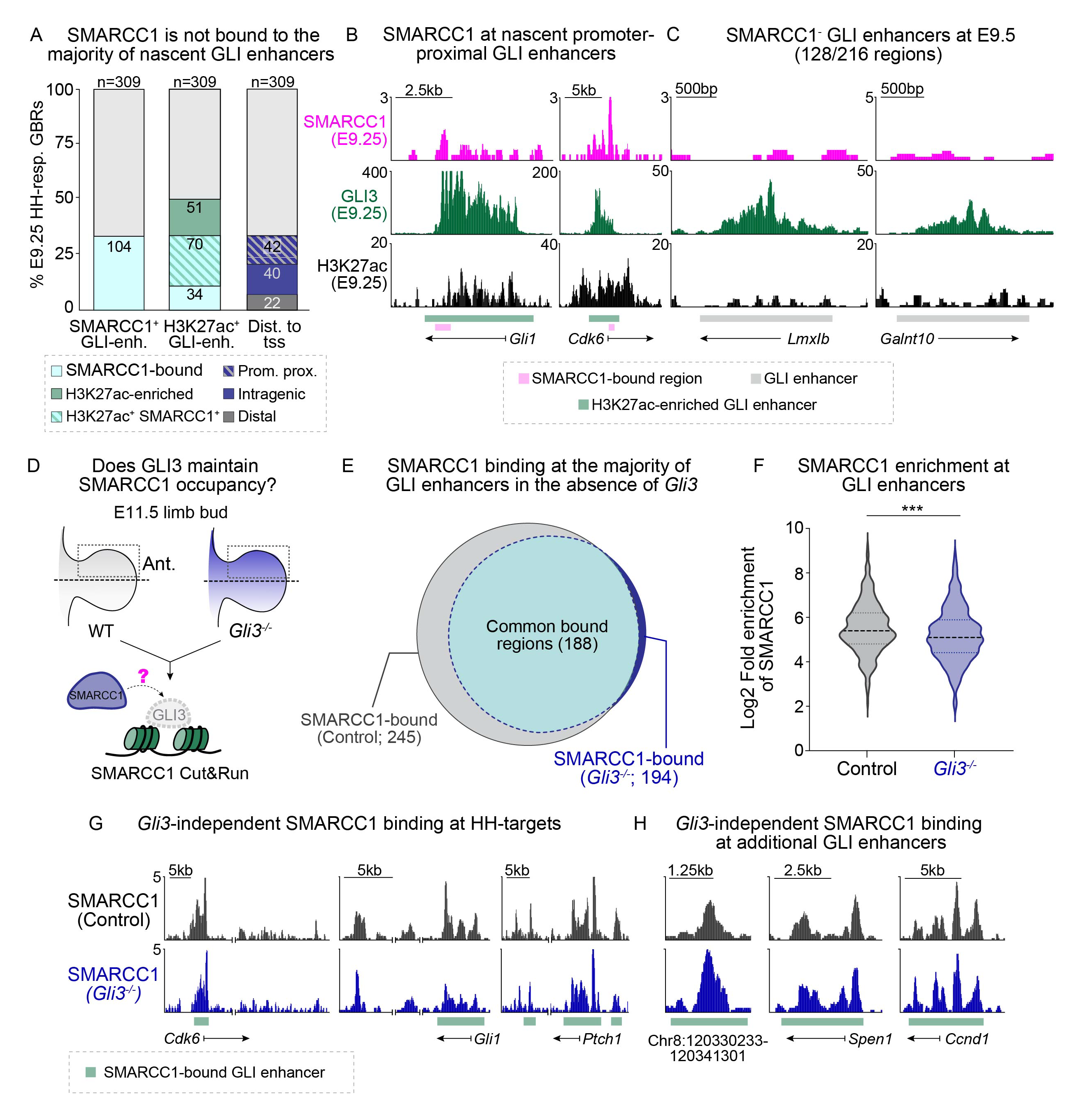
GLI3 does not recruit or maintain SMARCC1 at enhancers. A. SMARCC1 binds to a minority of H3K27ac-enriched (H3K27ac^+^), promoter-proximal GLI-bound enhancers (GLI-enh.) at E9.25 (21-24s). B,C. Representative tracks showing GLI3 and H3K27ac enrichment at previously defined nascent GLI enhancers with (B) and without (C) SMARCC1 binding (magenta) at E9.25. D. SMARCC1 binding determined by CUT&RUN on control (*Gli3^+/+^*; n=3) and *Gli3^-/-^*(n=4) anterior forelimbs at E11.5 (40-43s). E. Intersection of significant (FDR<0.05) SMARCC1 enrichment at GLI enhancers in control and *Gli3^-/-^*forelimbs. F. Fold enrichment of SMARCC1 at GLI enhancers in control and *Gli3^-/-^* forelimbs (Wilcoxon signed rank test). G,H. Representative tracks showing SMARCC1 binding at GLI enhancers.

To determine if GLI3 maintains SMARCC1 after the onset of HH signaling, we examined differential SMARCC1 binding in sibling control (*Gli3^+/+^*) and *Gli3^-/-^* anterior halves of forelimbs at E11.5, when SMARCC1 and GLI3 binding co-occur (Figure 3D). SMARCC1 was bound to 79% of GLI enhancers (245/309) in control limbs, and the majority of these regions (188/245) remained bound by SMARCC1 in *Gli3^-/-^* limbs (Figure 3E, Source data 4). This indicates that SMARCC1 is bound to most GLI enhancers even in the absence of *Gli3*, including biologically validated HH targets such as *Ptch1*, *Gli1* and *Cdk6* (Figure 3E, G-H). The 57 regions that appeared to lose SMARCC1 in *Gli3^-/-^*limbs generally had some detectable SMARCC1 binding in 1 or more mutant replicates, suggesting that variability among *Gli3^-/-^* embryos may have contributed to a loss in the number of significantly called SMARCC1 peaks (Figure 3E, Supp. Figure 3). Although SMARCC1 remained bound to most GLI enhancers in the absence of GLI3, there was a small but significant reduction in the overall enrichment levels of SMARCC1 in *Gli3 ^-/-^* limbs compared to controls for reasons that are presently unknown (Figure 3F, Source data 4). Overall, SMARCC1 is retained at the majority of GLI enhancers in the absence *Gli3*, suggesting that GLI3 likely does not recruit or maintain SMARCC1 at enhancers, but instead that SMARCC1 binds and regulates chromatin accessibility at enhancers in a GLI3-independent fashion. These results do not exclude a secondary role for a specific type of BAF complex in mediating later GLI-specific chromatin compaction after enhancer activation and given previous reports of *Smarcc1*-dependent de-repression of some HH-target genes in the limb (Jeon and Seong, 2016), we sought to determine if SMARCC1 is required for the transcription of HH target genes.

### *Smarcc1* is not required for most HH target gene expression

We compared gene expression between wild-type controls and SMARCC1 cKOs in E11.5 limb buds. The 928 differentially expressed genes (FDR<0.05) included approximately one-third (19/63) of previously predicted direct GLI target genes including *Gli1* and *Hhip* (which were both up-regulated in SMARCC1 cKO limbs) (Lex et al., 2022) (Supp. Figure 4A-H; Source data 5). In addition, if SMARCC1 mediates GLI3 repression, we reasoned that there would be concordance between genes upregulated in SMARCC1 cKOs and those upregulated in *Gli3^-/-^* limb buds.

**Figure 4.**
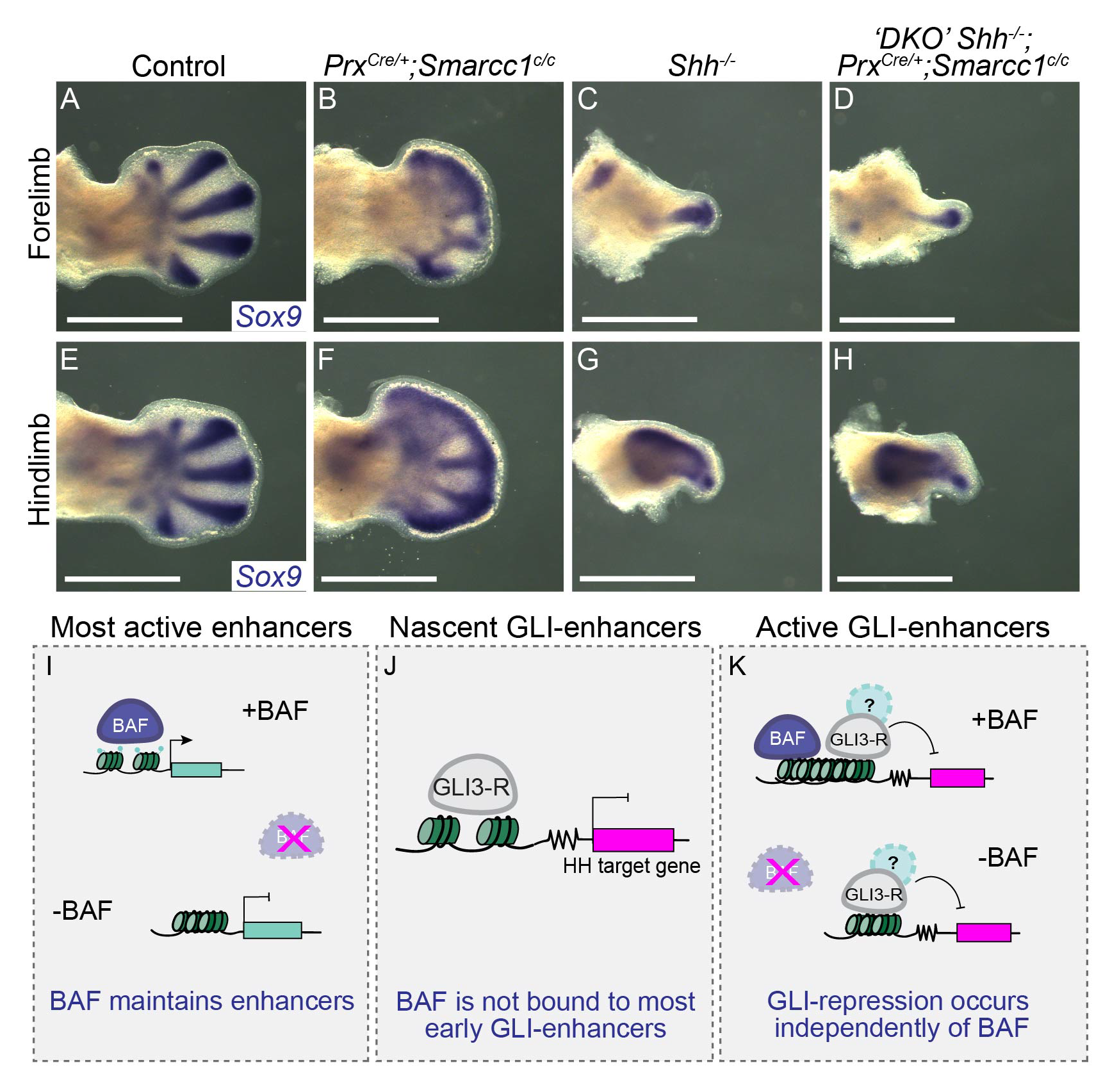
*Smarcc1* is not required for GLI-repression. A-H. *Sox9* expression in control (n=3; A,E), *PrxCre^+^;Smarcc1^c/c^* (n=4; B,F), *Shh^-/-^* (n=4; C,G), and *PrxCre^+^;Smarcc1^c/c^; Shh^-/-^* (‘DKO’) (n=3; D,H) fore-(A-D) and hindlimbs (E-H) at E12.5. Scale bars denote 10mm. I-K. Proposed model showing BAF-independent GLI transcriptional repression.

However, most GLI3-responsive genes previously shown to be upregulated in *Gli3^-/-^* limb buds (125/146; (Lex et al., 2022)) were not significantly changed in the absence of *Smarcc1.* The 21 *Gli3*-responsive genes that were changed were primarily downregulated (16/21) in SMARCC1 cKOs rather than upregulated (5) as would be expected if they were regulated by GLI3 repression (Supp. Figure 4B). This suggests that most GLI-repressor target genes are not SMARCC1-dependent. A caveat to this interpretation is that the *Hand2* expression domain, previously shown to be anteriorly expanded in *PrxCre^+^;Smarcc1^c/c^* limb buds (Jeon and Seong, 2016), was not upregulated in our experiments. This indicates that there are limitations to the sensitivity of the RNA-seq performed on whole limb buds that are likely leading to an under-reporting of GLI3-responsive genes.

To further examine potential direct repression targets of SMARCC1, we identified ATAC regions that were both significantly altered in SMARCC1 cKOs and bound by SMARCC1, and identified the nearest gene. We then intersected these genes with significantly up-and down-regulated genes identified by RNA-seq. Since enhancers that are repressed by GLI3 have reduced ATAC accessibility (Lex et al., 2022, 2020), enhancers that were co-repressed by SMARCC1 and GLI3 should have increased ATAC accessibility in SMARCC1 cKOs. Counter to this scenario, only 1/62 regions that had increased ATAC-accessibility in SMARCC1 cKOs was also bound by SMARCC1 (Source data 3). This region is nearest to *Prrx1*, which is down-regulated rather than upregulated in SMARCC1 cKOs. Indeed, many significantly down-regulated genes were bound by SMARCC1 and showed significant reductions in ATAC-accessibility in SMARCC1 cKOs (127/488), Overall, this suggests that SMARCC1 does not mediate the reduced chromatin accessibility mediated by GLI3 repression.

Pathway analysis of *Smarcc1*-responsive genes indicated enrichment of the WNT, Hippo, BMP and FGF signaling pathways as well as limb-specific transcription factors (i.e. *Tbx3*, *Tbx4*, and *Alx4*) (Supp. Figure 4A-H). Overall, this suggests that SMARCC1 regulates gene expression underlying many limb pathways but does not have a direct role in GLI transcriptional regulation. This is further supported by differences in the polydactylous phenotype observed in SMARCC1 cKOs compared to *Gli3^-/-^* limb buds. Consistent with prior reports, SMARCC1 cKOs have variable numbers of forelimb digits ranging from oligodactyly to polydactyly with polydactyly in the hindlimbs (n=3; Supp. Figure 4D-G) (Jeon and Seong, 2016). In contrast, *Gli3^-/-^* forelimbs have uniform forelimb polydactyly and lack the distinct distal digit fusions present in SMARCC1 cKOs mutants, suggesting that the digit phenotypes observed in SMARCC1 cKOs are likely not due to GLI de-repression alone (Bowers et al., 2012; Mo et al., 1997).

### SMARCC1 is not required for GLI3 repression

The above experiments suggested that SMARCC1 does not uniformly mediate global GLI repression. However, some HH target genes do require SMARCC1 for normal expression, suggesting that BAF complexes might still co-regulate a subset of GLI targets that are critical for limb development. *Shh;Gli3* double mutants have markedly improved digit and limb phenotypes compared to *Shh*^-/-^ embryos, indicating that loss of GLI3 repression is sufficient to rescue many aspects of the *Shh^-/-^* phenotype (Chiang et al., 2001; Lewandowski et al., 2015; Litingtung et al., 2002; teWelscher et al., 2002). If SMARCC1 is required for GLI3 repression, then the loss of *Smarcc1* should result in a partial rescue of digit number and limb size in *Shh^-/-^*limb buds. To test this, we compared the phenotypes in *Shh^-/-^* and *PrxCre^+^;Smarcc1^c/c^* (hereafter referred to as ‘DKO’) limb buds at E12.5, a stage when digit specification has already occurred. Wild-type embryos had 5 distinct *Sox9*-expressing digit condensates on both their forelimbs and hindlimbs (n=3) while *PrxCre^+^; Smarcc1^c/c^* limb buds had variable numbers of abnormally patterned condensates (n=4) (Figure 4A-B, E-F). In contrast, loss of *Smarcc1* was unable to rescue the *Shh* phenotype: DKO forelimbs and hindlimbs were indistinguishable from *Shh^-/-^* (n=4 of each genotype; Figure 4C-D, G-H). A limitation to interpreting these results is that GLI3 repression and HH signaling have essential early roles in limb specification and thus it is possible that *Smarcc1* conditional deletion occurs too late to rescue the phenotype (Lex et al., 2022; Scherz et al., 2007; Zhu et al., 2022, 2008). However, the conditional deletion of *Shh* results in a hypomorphic phenotype with reduced numbers of digits, suggesting that inhibition of GLI repression should have resulted in some improvement of the phenotype. In addition, the onset of *PrxCre* occurs earlier in the hindlimb than in the forelimb yet there was no improvement in the DKO hindlimbs either (Logan et al., 2002). Finally, SMARCC1 does not bind to the majority of GLI3-bound HH-responsive enhancers at early stages (Figure 3A, C) suggesting that it likely has a very limited role, if any, in GLI enhancer regulation prior to the onset of HH-signaling. Overall, this study shows that SMARCC1 does not directly repress the transcription of the majority of HH-target genes, does not require GLI3 for binding to chromatin, does not promote reductions in chromatin accessibility and is not genetically sufficient for mediating GLI3 repression. Our findings confirm previous reports that *Gli1* is ectopically expressed in *Smarcc1* mutant limb buds (Jeon and Seong, 2016). Ectopic *Gli1* has also been noted in *Smarcb1* mutant limb buds and in *Smarca4* mutant MEFs and neural cells, indicating it is generally regulated by BAF complexes (Jagani et al., 2010; Zhan et al., 2011). Under normal genetic conditions *Gli1* requires GLI-activator-dependent HH signaling and cannot be activated in *Shh;Gli3* double mutants that restore targets of GLI repression (Lewandowski et al., 2015; Litingtung et al., 2002; teWelscher et al., 2002). As *Ptch1* also requires GLI activation (Litingtung et al., 2002), the ectopic expression of *Ptch1* and *Gli1* in SMARCC1 cKOs cannot be explained by GLI de-repression. Our study does not explain why *Gremlin*, *Hand2* and *Hoxd13* are upregulated in *Smarcc1* mutant limb buds (Jeon and Seong, 2016). All three are negatively regulated by GLI3 repression but the positive regulators of their expression are not well understood (Lewandowski et al., 2015). We speculate that the misexpression of one or more of these positive factors in SMARCC1 cKOs causes their misexpression. SMARCC1 cKOs have widespread gene misexpression (929 differentially expressed genes, Source Data 5, Supp. Figure 4). This type of mis-regulation is also consistent with our finding that thousands of limb enhancers have altered chromatin accessibility in SMARCC1 cKOs.

Despite demonstrated binding to BAF complex members, including SMARCC1, GLI3 repressor functions independently of SMARCC1 at GLI enhancers. In the absence of a specific role in GLI repression, we propose that BAF complexes are independently enriched at most active enhancers, including GLI enhancers, where they regulate chromatin accessibility. Presumably, in this context they may also mediate context-specific aspects of GLI activation. This is distinct from parallel GLI transcriptional repression that recruits still unknown chromatin complexes to prevent gene expression (Figure 4I-K). These complexes either bind to GLI3 directly or indirectly to facilitate HDAC activity and chromatin accessibility at GLI enhancers (Lex and Vokes, 2022).Thus,

## METHODS

### Mouse strains and limb tissue isolation

Experiments involving mice were approved by the Institutional Animal Care and Use Committee at the University of Texas at Austin (protocol AUP-2022-00221). The *Smarcc1^tm2.1Rhs^* conditional allele (Jeon and Seong, 2016) (referred to as *Smarcc1^c/+^*) was maintained on a C57/BL6 background and crossed with *Prx1Cre^+^* mice (Logan et al., 2002) to generate control (*PrxCre^+^;Smarcc1^c/+^*) and *PrxCre^+^;Smarcc1^c/c^* embryos. The *Shh^tm1amc^* null allele (Dassule et al., 2000) (referred to as *Shh^+/−^*) was maintained on a Swiss Webster background and crossed with *PrxCre^+^;Smarcc1^c/+^* mice to generate double knockout (*PrxCre^+^;Smarcc1^c/c^;Shh^-/-^*; referred to as DKO) embryos. The *Gli3^Xt-J^* (Jackson Cat# 000026; referred to as *Gli3^+/-^*) allele was maintained on a Swiss-Webster background.

### *In situ* hybridization

For colorimetric in situ hybridization embryos were treated with 10µg/mL proteinase K (Invitrogen 25530015) for 30 minutes, post-fixed, pre-hybridized at 68°C for 1hour and hybridized with 500ng/mL digoxigenin-labeled riboprobes at 68°C overnight as previously described (Lewandowski et al., 2015). BM purple (Sigma-Aldrich 11442074001) staining was visualized in dissected forelimbs using a table-top Leica light microscope equipped with a DFC420C Leica camera.

Fluorescent in situ hybridization was conducted using HCR v3.0 reagents as previously described (Ramachandran et al 2022). Briefly embryos were fixed overnight at 4°C in 4% PFA/PBS and rehydrates with MeOH/PBST (0.1% Tween-20). Embryos were then treated with 10µg/mL proteinase K for 30 minutes, incubated in 4nM probe overnight at 37°C, and then in 60pmol hairpin overnight at room temperature. After hairpin incubation, the samples were washed and counterstained in DAPI (1:5000 dilution; Life Technologies D1306), embedded in low-melt agarose and cleared in CeD3++ as described (Anderson et al., 2020). Samples were imaged on a Nikon W1 spinning disk confocal.

### RNA-seq

Paired forelimbs were dissected from 2 individually genotyped control and *PrxCre^+^;Smarcc1^c/c^* embryos at E11.5 (39-40 somites). RNA was extracted using Trizol reagent (Life Technologies, 15596026). Libraries were generated by BGI Genomics: samples were 3’adenylated, adaptor ligated, UDG digested, and circularized following amplification to generate a DNA nanoball that was then paired-end sequenced on a DNBSEQ platform at a depth of 40 million reads/sample. Sequenced reads were aligned to the mm10 mouse genome using Salmon (version 1.9.0) and assembled into genes using R package tximeta (version 1.16.0) (Love et al., 2020). Differential gene expression was calculated using DESeq2 (Love et al., 2014). We defined significantly differentially expressed genes using an FDR<0.05. Differentially expressed genes are listed in Source data 5.

### ATAC-seq

Anterior forelimb pairs from 2 E11.5 (40-48s) control (*PrxCre^-^;Smarcc1^c/c^*) and 3 *PrxCre^+^;Smarcc1^c/c^* embryos were dissected and each was used as a single biological replicate. ATAC-Seq was performed as described previously (Lex et al., 2022). Briefly limbs were dissociated in 100µg/mL Liberase (Roche 05401119001), lysed for 10minutes at 4°C in nuclear permeabilization buffer and centrifuged for 10 minutes. Nuclei were then resuspended, and 50,000 nuclei from each replicate were incubated with 2.5µl Transposase (Illumina 20034210) in a 50µl reaction. The transposase reaction was carried out for 1hour at 37°C, shaking gently. Libraries were generated using the NEB High-Fidelity 2x Mix (New England Biolabs M0541S), PCR amplified for 11 cycles and cleaned up using SparQ PureMag beads (QuantaBio 95196- 060). Libraries were sequenced using paired-end 75bp reads at a depth of ∼40 million reads/sample. Samples were mapped to the mm10 mouse genome using Bowtie2 (version 2.3.5) (Trapnell et al., 2012), peaks were called using MACS2 (Zhang et al., 2008), normalized using library size parameters, and differential analysis was performed using Limma (Ritchie et al., 2015). Significantly changed peaks were called using an FDR<0.05. Differentially ATAC-accessible regions are listed in Source data 3. To identify direct targets of SMARCC1, significantly ATAC-changed regions (62 with increased ATAC accessibility and 10,844 regions with reduced ATAC accessibility) were intersected with SMARCC1-binding regions in the wild-type anterior limb bud. Nearest gene analysis was performed using GREAT (using a cutoff of +/-500KB) on SMARCC1-bound, ATAC-changed regions, and the resulting gene list was intersected with genes that were significantly up-regulated and down-regulated in SMARCC1 cKOs. The intersections for this analysis are provided in Source data 3.

### Chromatin binding

For wild-type (Swiss Webster) CUT&RUN experiments, pairs of whole forelimbs at E9.25 (21-24 somites) were pooled to allow for sufficient cell numbers; for E11.5 (43-44 somites), anterior and posterior halves of forelimbs were processed individually. 2 biological replicates were collected at E9.25 and 3 biological replicates were collected at E11.5. For differential CUT&RUN, the anterior halves of forelimb pairs were collected from single, genotyped control (*Gli3^+/+^)* and *Gli3^-/-^* embryos. 3 biological replicates were collected for each genotype. Dissected tissues were dissociated in 100µg/mL Liberase at 37°C for 10minutes and processed using EpiCypher’s CUTANA kit reagents (EpiCypher 15-1016) and protocol with the following modifications. Following dissociation, samples were incubated overnight at 4°C with 1:50 SMARCC1 primary antibody (Invitrogen PA5-30174). Samples were then incubated with 1:100 donkey anti-rabbit (Jackson ImmunoResearch 711-035-152) secondary antibody for 30 minutes at room-temperature, washed in digitonin wash buffer and incubated with CUTANA pAG-MNase for 10minutes at RT. The MNase reaction was then performed at 4°C for 2 hours. Libraries were generated using the NEBNext Ultra II DNA library prep kit (New England Biolabs E7645S) with 14 PCR cycles using unique dual indexes (New England Biolabs E6440S). Following clean-up with SparQ PureMag beads (QuantaBio), samples were sequenced using PE150 at a read depth of ∼5 million reads. Samples were mapped to the mm10 mouse genome using Bowtie2 (version 2.3.5) (Trapnell et al., 2012), peaks were called using MACS2 (Zhang et al., 2008). Called peaks for wild-type CUT&RUN for SMARCC1 at E11.5 and E9.25 and a list of overlapping VISTA enhancers at E11.5 are provided in Source data 1.

Differential SMARCC1 binding between E11.5 (40-43s) control (*Gli3^+/+^*; n=3) and *Gli3^-/-^* (n=4) samples was conducted using an E.coli spike-in (EpiCypher 18-1401) of 0.125ng/sample. Samples were sequenced on the Novaseq platform using PE150 at a read depth of ∼7 million reads. Sequenced reads were aligned to the mouse reference genome mm10 using Bowtie2 (Trapnell et al., 2012) and peaks were called using MACS2 (Zhang et al., 2008) with an FDR<0.05. For Figure 3E, enrichment values were counted from the previously published list of 309 GLI enhancers (Lex et al., 2022) and normalized using library size first, and then were Log2 transformed. For control and mutant groups respectively, the average value across biological replicates was used to calculate the final fold enrichment. All called peaks are listed in Source data 4.

Previously published H3K27ac ChIP-seq samples from E11.5 forelimbs (n=2) (GEO accession # GSE151488; (Malkmus et al., 2021)) were aligned to the mm10 mouse genome to generate ChIP-seq tracks and peak-called using Bowtie2 (Trapnell et al., 2012) and MACS2 (Zhang et al., 2008) respectively with an FDR<0.05 as cutoff. Significantly called peaks are listed in Source data 1.

### Western Blots

Western blots were performed on individually genotyped E11.5 (42-45s) *PrxCre^-^*;*Smarcc1^c/+^*(control, 3 replicates) and *PrxCre^+^;Smarcc1^c/c^* (3 replicates) forelimb pairs. Limbs were lysed in 80µl of RIPA buffer (10mM Tris pH 7.4, 150mM NaCl, 1mM EDTA pH 8, 1% sodium Deoxycholate (wt/vol), 0.1% SDS (wt/vol), 1% Triton X-100 (vol/vol)) and 10µl of each sample was loaded onto a 4-20% gradient agarose gel, blocked in 5% BSA for 1 hour at room temperature, and then incubated with the following primary antibodies for 2 hours at room temperature in 5% BSA: 1:1000 anti-SMARCC1 (Cell Signaling Technologies cat. #11956T), 1:1000 anti-SMARCA4 (Cell Signaling Technologies cat. #49360T), 1:1000 anti-SMARCE1 (Cell Signaling Technologies cat. #33360T), 1:1000 anti-SMARCC2 (Cell Signaling Technologies Cat. #12760T), 1:1000 anti-ARID1A (Cell Signaling Technologies cat. #12354T), 1:1000 anti-ARID1B (Cell Signaling Technologies cat. #92964T) 1:2000 anti-GAPDH (Cell Signaling Technologies cat. #5174S). Samples were then incubated in secondary antibodies for 1 hour at room temperature in 3% BSA: 1:5000 Donkey anti-Rabbit (Jackson 711-035-152). Blots were developed using Amersham ECL Prim (Cytiva RPN2232). Protein expression levels were quantified by determining the band intensity in ImageJ using the gel analysis function. The intensities for the bands of the BAF subunit in each replicate were normalized by taking the ratio of the intensity for the band to the intensity of the corresponding GAPDH band. Uncropped Western blot images are provided in Source data 2.

### Skeletal Preparation

Skeletons of Postnatal (P) day 0 mouse pups were prepared as described (Allen et al., 2011). Briefly pups were harvested, skinned, eviscerated and fixed in ethanol and acetone. Skeletons were then stained in an Alcian blue/Alizarin red stain solution for 4 days, incubated in 1% potassium hydroxide and cleared in glycerol. Skeletons were imaged using a table top Leica light microscope.

## ACKNOWLEDGEMENTS

We thank Dr. Kyunghee Choi and Dr. Rho Hyun Seong for providing the *Smarcc1* mouse strain. We thank the Genomic Sequencing and Analysis Facility at UT Austin, Center for Biomedical Research Support (RRID#: SCR_021713). This work was supported by NIH R01HD073151 (to SAV and HJ) and F31DE027597 (to R.K.L.).

## SUPPLEMENTAL FIGURE LEGENDS

**Supplemental Figure 1.**
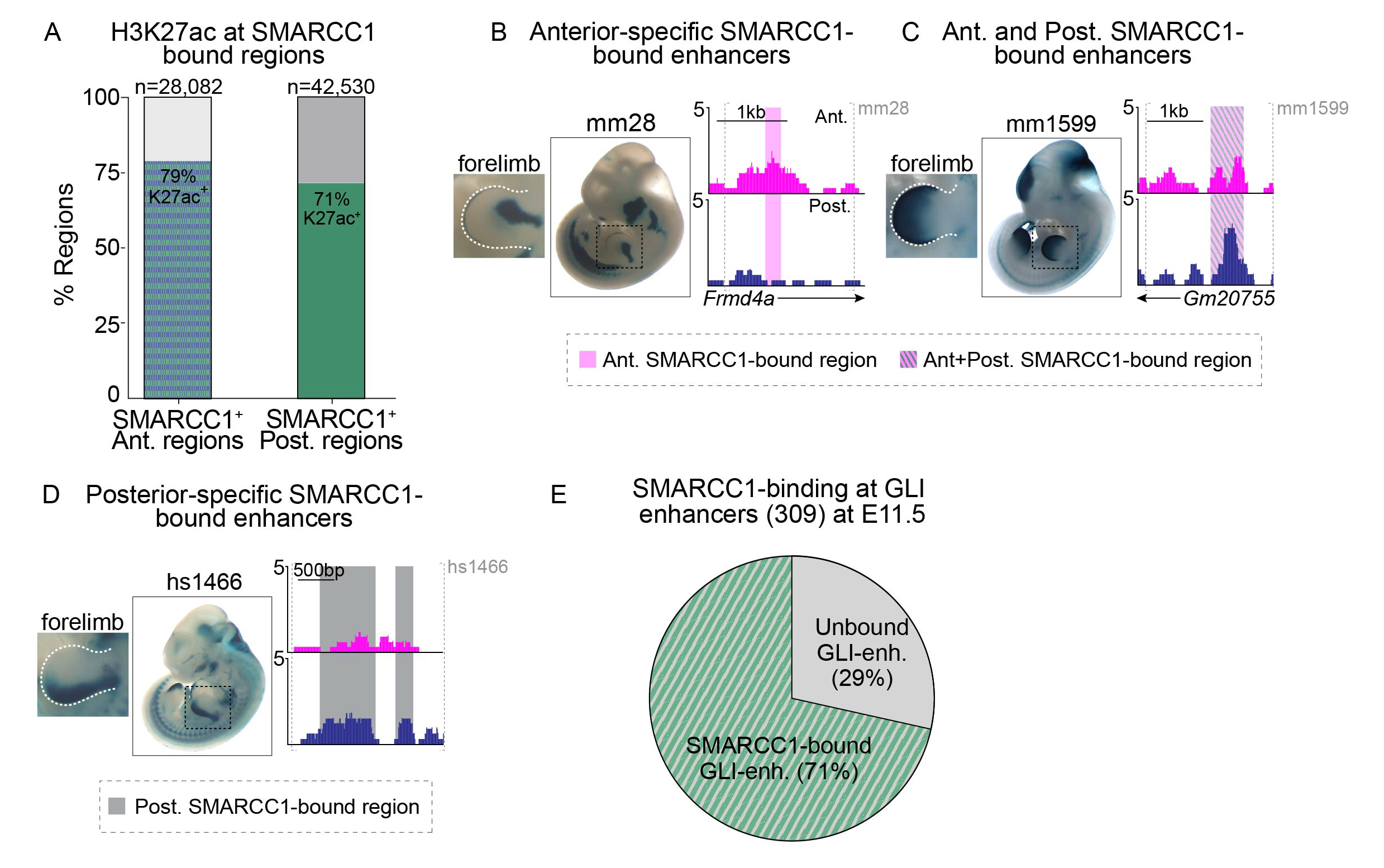
SMARCC1 is recruited to some region-specific limb enhancers and most GLI enhancers. A. H3K27ac enrichment at SMARCC1-bound regions in anterior and posterior limb buds. B-D. SMARCC1-bound limb VISTA enhancer regions with anterior-specific (B), broad (C), and posterior-specific (D) enhancer activity. E. SMARCC1 binding at GLI enhancers at E11.5.

**Supplemental Figure 2.**
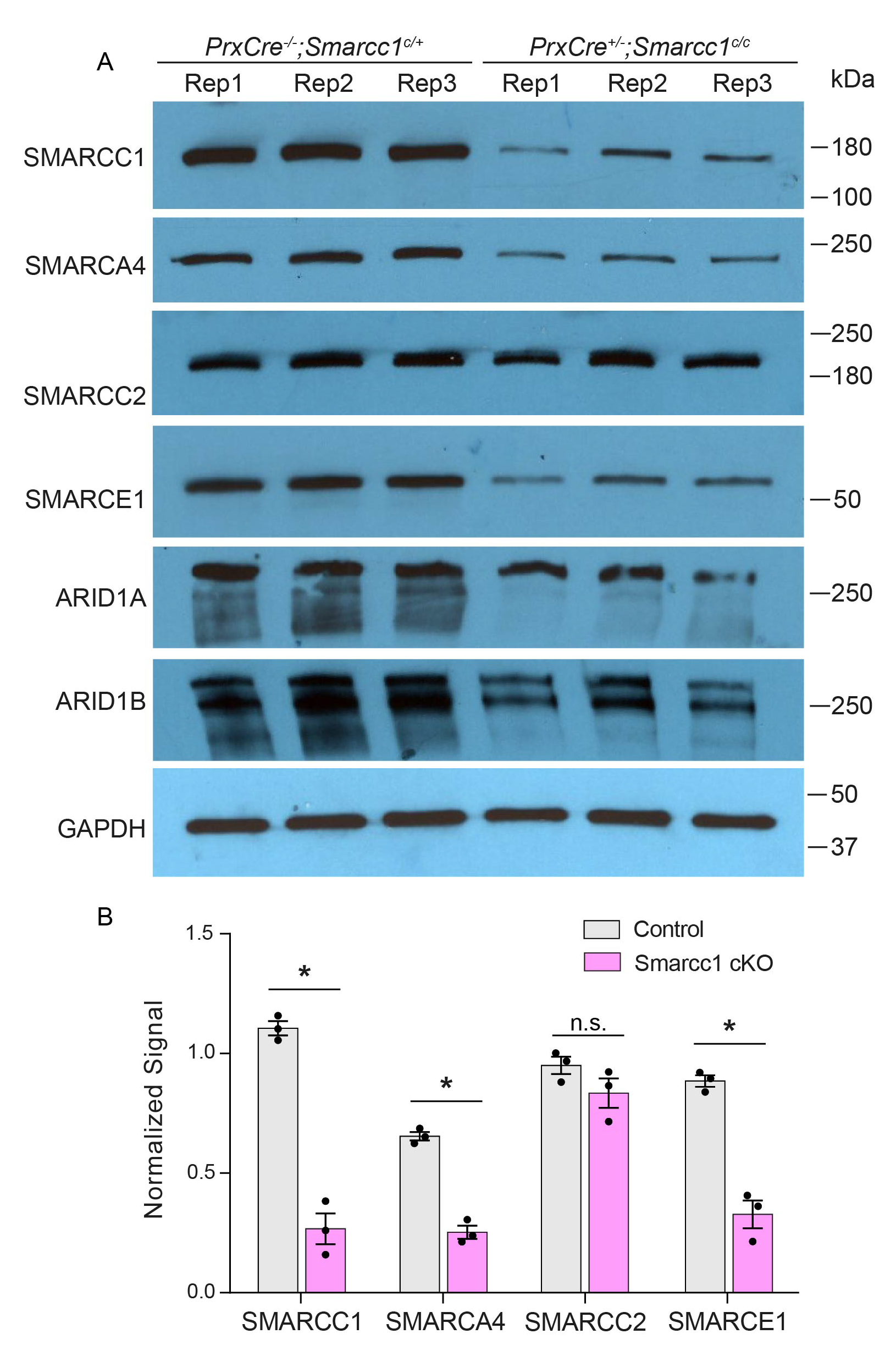
Several BAF complex members have reduced expression in SMARCC1 CKOs. A. Western blot showing expression of BAF complex members and GAPDH in forelimbs pairs from individual E11.5 control (*PrxCre*^-^;*Smarcc1*^c/+^) and SMARCC1 cKO (*PrxCre*^+^;*Smarcc*1^c/c^) embryos (three biological replicates). B) Quantification of changes in expression for select BAF subunits normalized to GAPDH. Note that despite appearing reduced, ARID1A/B were not quantified due to the presence of multiple bands. Error bars indicate standard error of mean (*p<0.001, Student’s T-test; n.s. denotes protein levels that are not significantly changed).

**Supplemental Figure 3.**
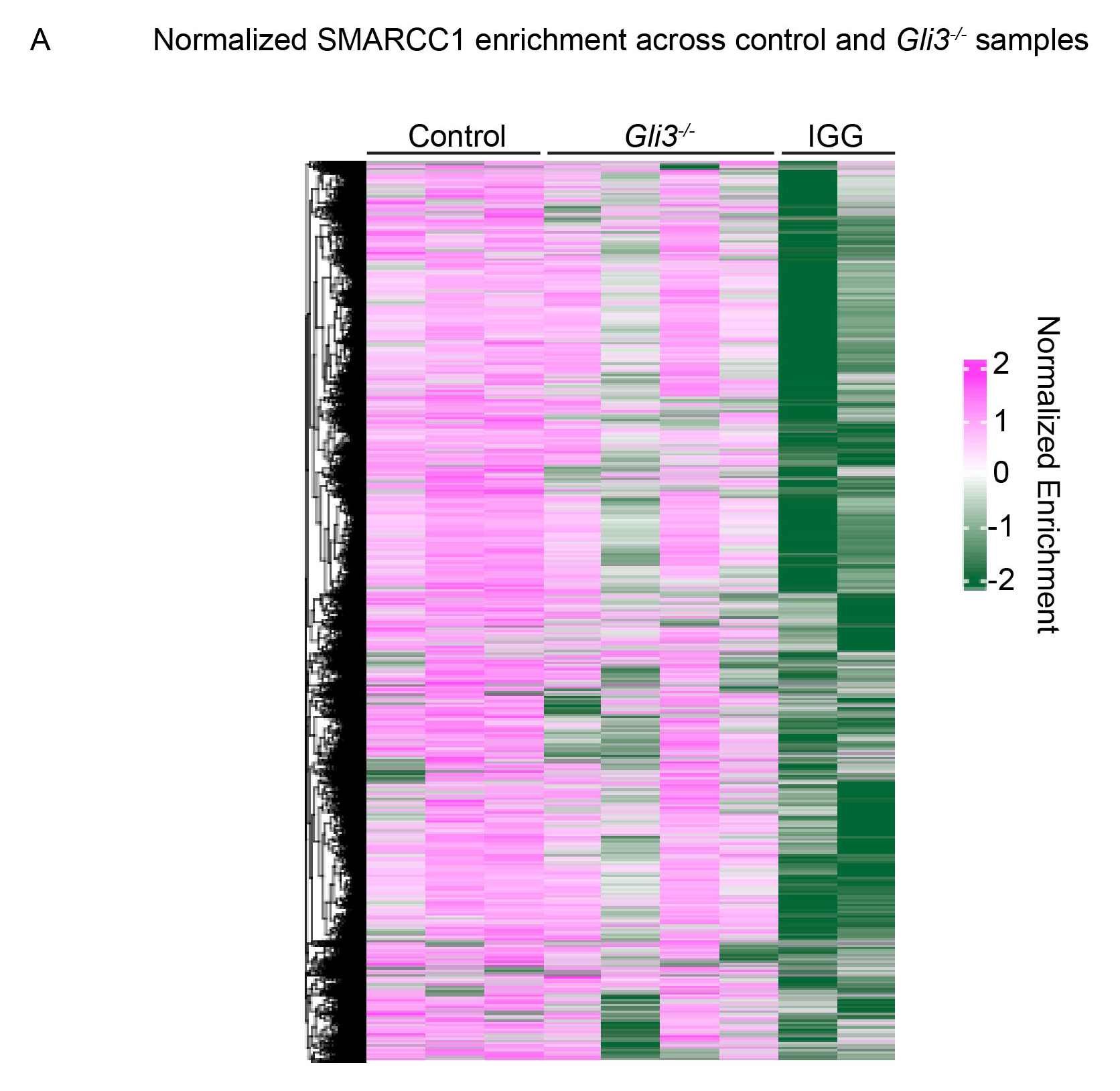
Sample variability in SMARCC1 binding within *Gli3^-/-^* limbs. A. Heatmap showing normalized genome-wide SMARCC1 enrichment within controls (*Gli3 ^+/+^*; n=3) and *Gli3^-/-^* limbs (n=4).

**Supplemental Figure 4.**
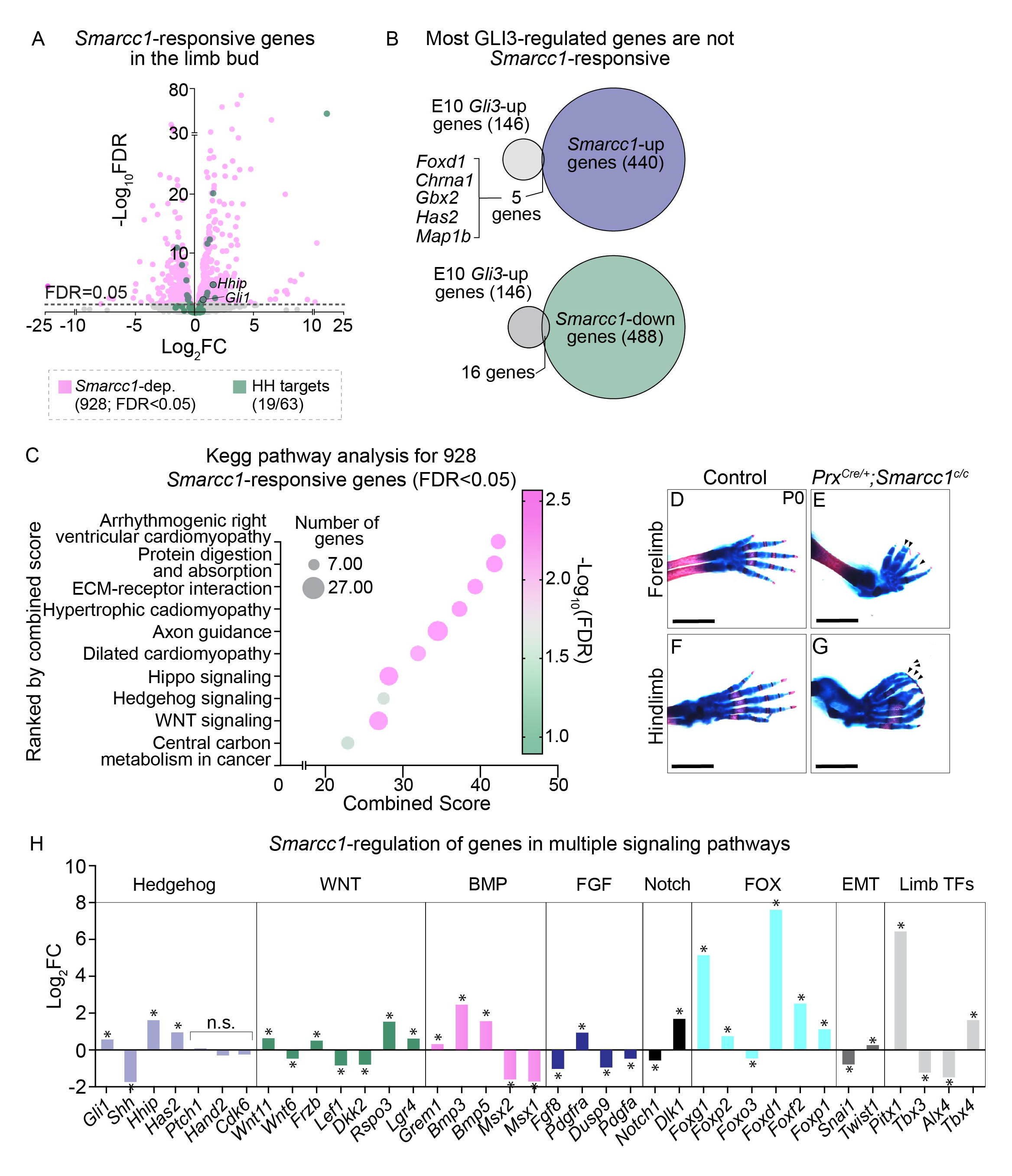
Loss of *Smarcc1* results in widespread disruption of signaling pathways in the limb. A. RNA-seq of E11.5 control and *PrxCre^+^;Smarcc1^c/c^* forelimbs (n=2 per condition). Significantly changed genes are marked in light pink and previously predicted HH-target genes (listed in Source data 4) are highlighted in green. B. Intersection of previously identified *Gli3*-upregulated genes (GEO accession # GSE178838) with either *Smarcc1*- upregulated or downregulated genes. C. KEGG Pathway analysis of 928 significant *Smarcc1*- responsive genes (FDR<0.05). D-G. Skeletal preparations of control (n=3) and *PrxCre^+^;Smarcc1^c/c^* (n=3) forelimbs (D-E) and hindlimbs (F-G) at P0. Scale bars denote 2mm. H. Fold enrichment of HH, WNT, BMP, FGF, Notch, FOX, EMT and Limb transcription factors (TFs). *p<0.05; n.s. denotes genes that are not significantly changed (p>0.05).

